# PAT mRNA decapping factors function specifically and redundantly during development in *Arabidopsis*

**DOI:** 10.1101/2022.07.06.498930

**Authors:** Zhangli Zuo, Milena Edna Roux, Yasin F. Dagdas, Eleazar Rodriguez, Morten Petersen

**Affiliations:** Department of Biology, Faculty of Science, University of Copenhagen, Copenhagen, Denmark; Gregor Mendel Institute, Austrian Academy of Sciences, Vienna BioCenter, Vienna, Austria

## Abstract

Evolutionarily conserved PAT1 proteins activate mRNA decay through binding mRNA and recruiting decapping enzymes and other factors hence optimize transcriptional reprogramming during development. Here, we generated multiple mutants of *pat1* (Protein Associated with Topoisomerase II), *path1* and *path2* and inspected their growth and leaf morphology phenotype. *pat* triple mutants exhibit extreme stunted growth and all mutants with *pat1* exhibit leaf serration while mutants with *pat1* and *path1* all display short petioles. All 3 PATs can be found localized to Prossessing Bodies (PBs) upon auxin treatment and RNA-seq analysis indicate that all 3 PATs redundantly regulate auxin responses. Moreover, shade avoidance and NAC genes are misregulated in *pat1path1* double and *pat* triple mutants suggesting PAT1 and PATH1 function in petiole elongation and leaf patterning. In conclusion, PAT proteins exhibit both specific and overlapping functions during different stages of plant growth and our observations underpin the importance of the mRNA decay machinery for proper development.

## Introduction

The RNA-binding proteins (RBPs) PAT1(Protein Associated with Topoisomerase II) family proteins are highly conserved through eukaryotes and mediate mRNA decay in cytoplasm. As a central platform, PAT1 forms a heterooctameric complex with LSM(Like-sm)1-7 at 3’ end of oligoadenylated mRNAs to engage transcripts containing deadenylated tails thereafter recruits decapping enzymes and other factors for 5’-3’ decay, these decapping complex and mRNAs can diffuse in cytoplast or aggregate into distinct cytoplasmic foci called PBs (Brengues et al., 2005; Balagopal and Parker, 2009;Ozgur et al., 2010; Chowdhury et al., 2014; Charenton et al., 2017; Lobel et al., 2019). The deadenylated mRNAs could also be degraded 3’-5’ via exosomal exonucleases and SUPPRESSOR OF VCS (SOV)/DIS3L2. mRNA decay regulates transcriptome shift and thereby impacts organism development and responses to stresses (Newman et al., 2017; Essig et al., 2018; Xu and Chua, 2012; Merret et al., 2013; Roux et al., 2015; Perea-Resa et al., 2016).

Yeast, fungi and green algae encode one PAT1 paralog respectively and vertebrates encode two while most plants possess multiple PAT1 family members (Zhang et al., 2020). For example, moss has two PAT1 paralogues, rice holds four and Arabidopsis encodes three, named PAT1, PATH1 and PATH2 (Zhang et al., 2020; Roux et al., 2015). Deletion of well conserved *PAT1* gene exhibits temperature sensitive phenotype in yeast and embryonic lethality in *Drosophila melanogaster* respectively (Wyers et al., 2000; Pradhan et al., 2012). Besides, we previously reported that Arabidopsis *pat1* mutant exhibit auto-immune phenotype, PAT1 (AtPAT1) localizes to PBs and is phosphorylated on serine 208 by MAP kinase 4 (MPK4) in response to bacterial flagellin (Roux et al., 2015). Loss-of-function of AtPAT1 inappropriately triggers the immune receptor SUMM2, and *Atpat1* mutants consequently exhibit dwarfism and de-repressed immunity (Petersen et al., 2000, Roux et al., 2015, Zhang et al., 2012). Thus, PAT1 is under immune surveillance and PAT1 function(s) are best studied in SUMM2 loss-of-function backgrounds (Roux et al., 2015).

In addition to PAT1, Arabidopsis encodes 2 other PATs, PATH1 and PATH2. So far, little is known about the functions of these 3 PATs during plant development. Here we have disrupted the mRNA decapping components PAT1, PATH1 and PATH2 in the *summ2* background (Roux et al., 2015) which allows us to study this process without autoimmunity interference. We have revealed all three PAT proteins interact with the LSM1 decapping factor and localize to PBs upon different stimuli perception. Through observing developmental phenotype of multi-*pats* mutants and performing RNA-seq to examine differently expressed genes we identified PAT1 is mainly responsible to keep regular leaf pattern and work redundantly with other 2 PATs during plant development.

## Results

### PAT mRNA decapping factors interact with LSM1

Arabidopsis genome encodes 3 PAT mRNA decapping factors: PAT1, PATH1 and PATH2 (Roux et al., 2015). We previously reported that PAT1 localizes to PBs and can be found in complexes together with the decapping component LSM1 (Roux et al., 2015). To test if PATH1 and PATH2 are also in mRNA decay complex, we transiently expressed all three PAT-HA fusions with YFP-LSM1 and performed coimmunoprecipitation assays. All three PAT-HA fusions could be detected in LSM1 immunoprecipitates (Fig 1A), indicating that, similar to PAT1, PATH1 and PATH2 also localize to LSM1 positive compartments. In line with this, a previous study looking for LSM1 associated proteins detected all three PATs proteins in LSM1 pull downs (Golisz et al., 2013).

**Fig 1.**
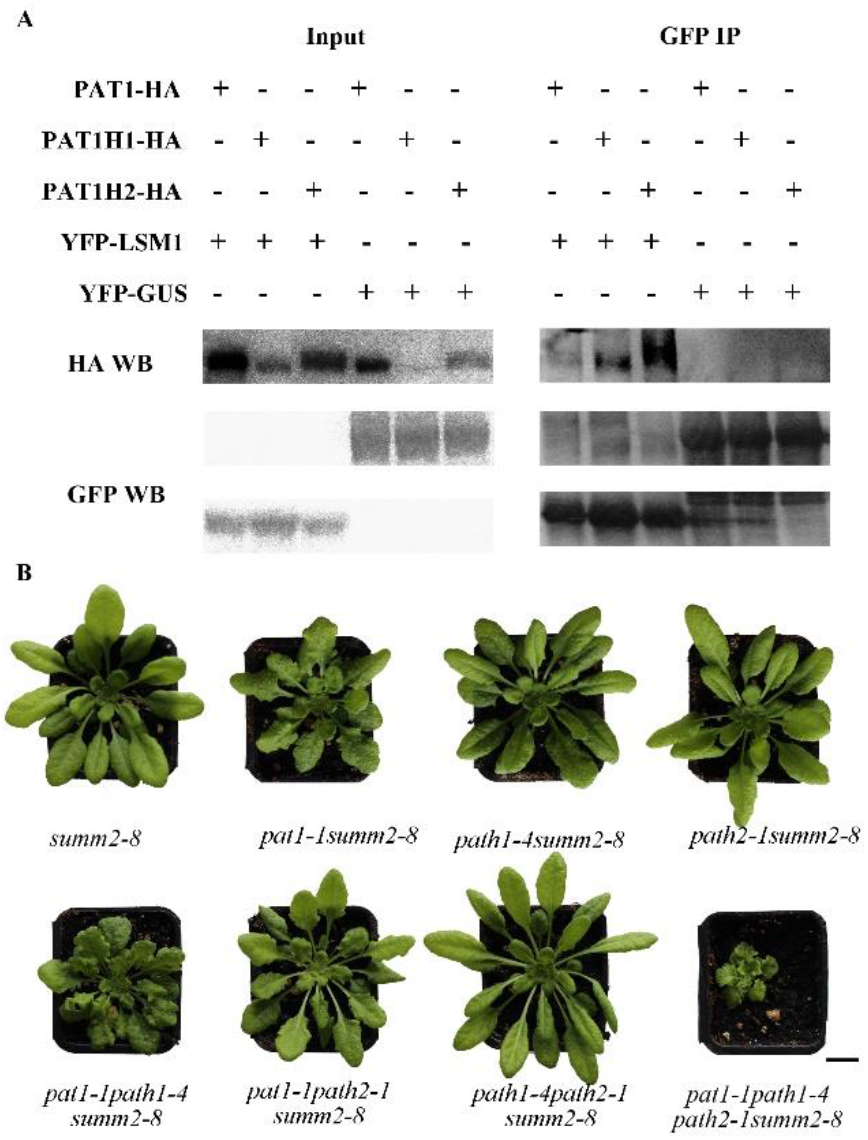
PATH1 and PATH2 are also mRNA decapping factors. (**A**) Co-IP between the three PAT-HA and YFP-LSM1 fusions. Proteins were transiently co-expressed in *N. benthamiana* and tissue harvested 3 days post-infiltration. Immunoblots of inputs (left panels) and GFP IPs (right panels) were probed with anti-HA antibodies (upper pannel) and anti-GFP antibodies (middle panel for YFP-GUS and bottom panel for YFP-LSM1). (**B**) 6-week-old plants of *summ2-8* and multiple *pat* mutants grown in soil in a chamber with 8/16hrs light/dark at 21°C. A representative plant for each line is shown. The scale bar indicates 1cm.

### PAT(s) loss-of-function mutants exhibit different growth phenotypes

While we had detected PAT1 in PBs and found it under SUMM2 surveillance (Roux et al., 2015), little was known about mRNA targets or functions of the three PATs in *Arabidopsis*. To address this, we generated single, double and triple knockouts (KOs) of the three PATs in the *summ2-8* background to avoid immune activation. *path1* and *path2* mutants were therefore generated in *pat1-1summ2-8* and *summ2-8* by CRISPR/CAS9-mediated genome editing (Fig 1B&S1A). Fig S1B shows the sequences of *PATH1* and *PATH2* in 2 independent *pat* triple mutants. The growth phenotypes of 6 weeks-old soil-grown plants of *pat* single, double and triple mutants at 21 °C with 8/16 hrs light/dark photoperiod are shown in Fig. 1B& S1A. Both independent lines of *pat1-1path1-4path2-1summ2-8* and *pat1-1path1-5path2-2summ2-8* exhibited markedly stunted growth compared to the other *pat* single or double mutants (Fig. 1B, S1B) indicating functional redundancy of PATs in regulating plant development. *path1-4summ2-8* and *path2-1summ2-8* have leaf number and shape similar to *summ2-8* while *pat1-1summ2-8, pat1-1path1-4summ2-8, pat1-1path2-1summ2-8* and *pat1-1path1-4path2-1summ2-8* grow fewer leaves and with serration shape indicating PAT1 has specific function in regulating leaf serration (Fig. 1B). Furthermore, complementation lines of *pat1-1path1-4summ2-8* and *pat1-1path2-1summ2-8* with *Venus-PATH1* and *Venus-PATH2* respectively show the same growth phenotype as *pat1-1summ2-8* (Fig. S1C&D).

### PATs localize to PBs upon different stimuli perception

mRNA decapping complex and mRNAs can aggregate into PBs in the cytoplasm (Ozgur et al., 2010). To investigate if all three PAT proteins may also localize into PBs, we exposed seedlings expressing Venus-fusions of the PAT proteins (Fig. S1C-F) to the phytohormones auxin (IAA), cytokinin, ethylene precursor ACC, bacterial flagellin peptide flg22 and extracellular ATP. All stimuli induced massive increase of Venus-PAT1 distinct foci in roots within 20 min (Fig. 2). PATH1 only localized into PBs in response to IAA while PATH2 also enters PBs under ACC treatment (Fig 2). These observations indicate PAT1 plays a main role in forming PBs compared to PATH1 and PATH2.

**Fig 2.**
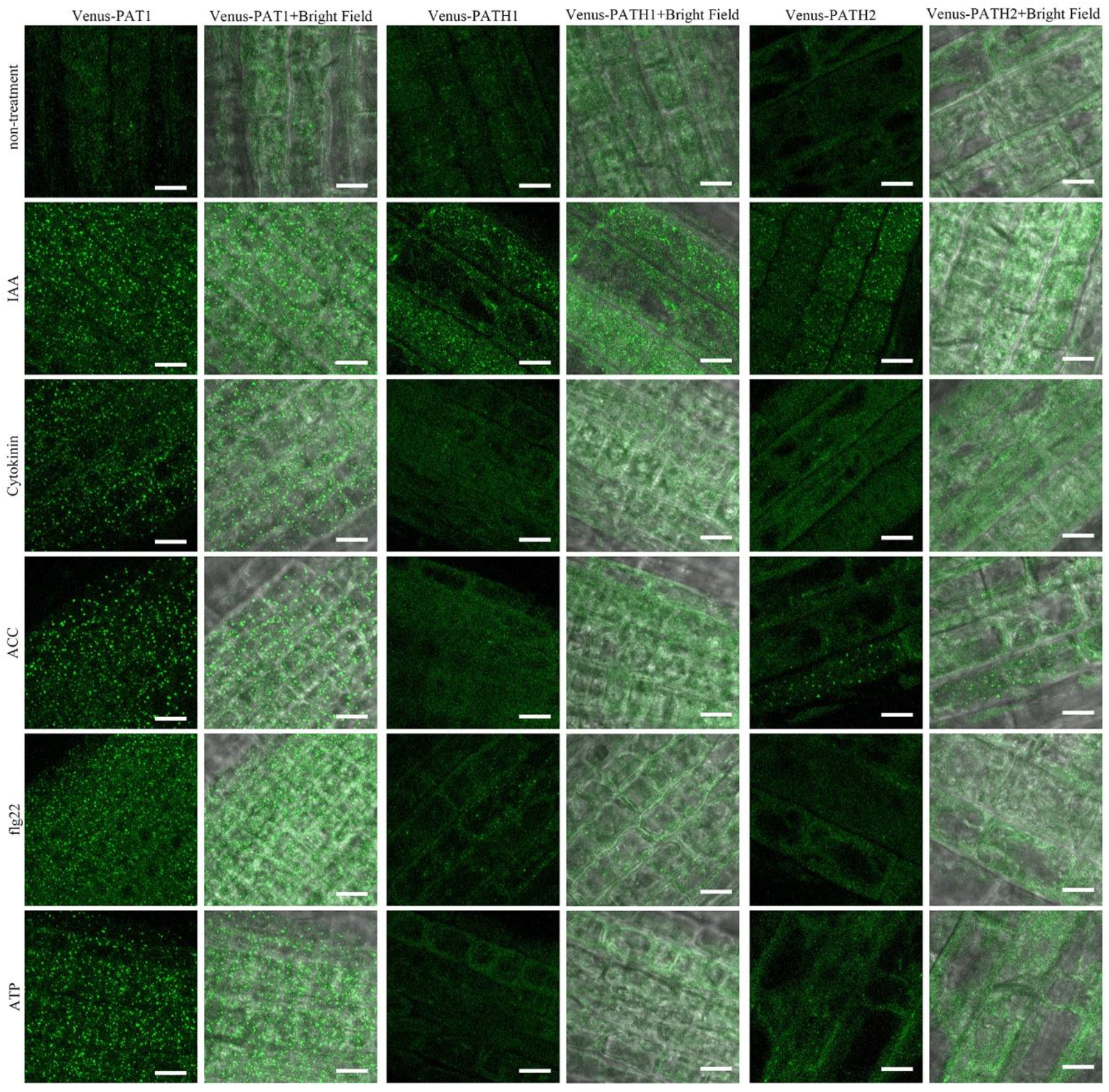
PATs relocalize to PBs upon stimuli perception. Venus-PATs expressing seedlings in MS growth medium or 20min after treatment with MS containing 0.5µM IAA, 1µM Cytokinin, 25µM ACC, 5µM flg22 or 0.1mM ATP. Representative images are shown. Scale bars indicate 10µm.

### All three PATs redundantly regulate plant development

To better quantify growth phenotype of these *pats* mutants, we dissected 6-week-old plants of Col-0 and multiple *pat* mutants (Fig 3A) and recorded leaf (leaf blade length >1mm) numbers. All mutants except *path1-4summ2-8* and *path2-1summ2-8* developed fewer leaves than Col-0 and *summ2-8*, while *pat1-1path1-4path2-1summ2-8* exhibit least leaf numbers (Fig 3B), indicating redundancy of all 3 PATs in regulating plant growth.

**Fig 3.**
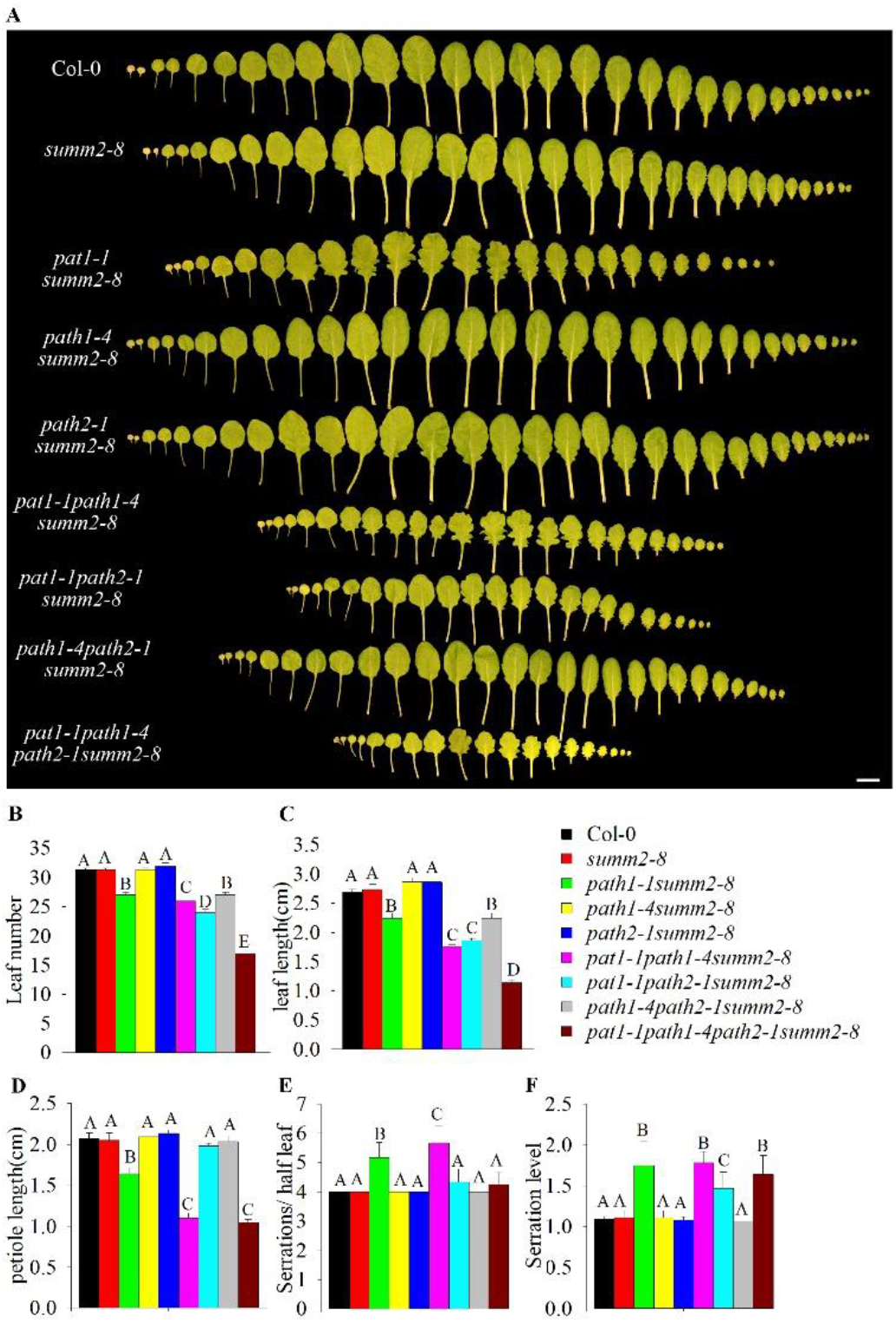
Leaf morphology of *pat* mutants. Leaf development (**A**), leaf numbers (**B**), leaf blade length (**C**), leaf petiole length(**D**), leaf serration numbers(**E**) and serration levels (**F**) of 6 week-old plants of Col-0, *summ2-8, pat1-1summ2-8, path1-4summ2-8, path2-1summ2-8, pat1-1path1-4summ2-8, pat1-1path2-1summ2-8, path1-4path2-1summ2-8* and *pat1-1summ2-8, path1-4summ2-8, path2-1summ2-8, pat1-1path1-4summ2-8, pat1-1path2-1summ2-8* and *pat1-1path1-4path2-1summ2-8*. The scale bar indicates 1cm. Leaf-serration levels are expressed as the distance from tip-to-midvein divided by the distance from sinus-to-midvein for different teeth.

### PATs play different roles in regulating leaf morphology

Leaf morphology is an essential part of phytomorphology. We therefor inspected leaf size and shape of the latest expanded leaves from the multiple *pat* mutants. Similar to leaf number, *pat1-1path1-4path2-1summ2-8* exhibit shortest leaf blade and all mutants except *path1-4summ2-8* and *path2-1summ2-8* developed shorter leaf blade than Col-0 and *summ2-8* (Fig 3C). Petioles connect leaf lamina to plant stem, *pat1-1summ2-8* exhibit shorter petioles than Col-0 and *summ2-8*, while *pat1-1path1-4summ2-8* and *pat1-1path1-4path2-1summ2-8* showed shortest petioles (Fig 3D), indicating only PAT1 and PATH1 are regulating leaf petiole length and PAT1 plays the main role.

Evolutionarily, variation in plant leaf shape reflects natural selection operating on function. To better understand PATs function in leaf serration, we counted serration numbers per half leaf and measured serration level on the latest expanded leaves from the multiple *pat* mutants (van Wijk et al., 2018). Interestingly, *pat1-1summ2-8* displayed increased number of serrations compared to control plant and *pat1-1path1-4summ2-8* but not *pat1-1path2-1summ2-8* showed the most serrations (Fig 3E), suggesting both PAT1 and PATH1 but not PATH2 function in regulating leaf serration numbers. All mutants with *pat1* loss-of-function exhibit higher serration level than other mutant combinations and *pat1-1path2-1summ2-8* displayed lowest serration level in plants with *PAT1* mutated (Fig 3F), indicating PAT1 is regulating leaf serration level, and PATH2 may play opposite role. All these results suggest that the function of PATH1 and PATH2 are redundant to the other PATs, while PAT1 serves a main and specific function in development.

### PAT(s) loss-of-function mutants exhibit different transcriptome

Plant and organ growth is tightly regulated by their developmental program. To identify genes which affect different developmental programs regulated by mRNA decapping factor PATs, we performed RNA-seq from plants of *pat1-1path1-4path2-1summ2-8* and all single and double mutant combinations shown in Fig 1B. Principal Component Analysis (PCA) Plot (Fig. 4A) exhibit general similarities of these 8 lines, revealing that mutants with *pat1* are different from the other plants based on PC1, and based on PC3, *pat1-1summ2-8* and *pat1-1path1-4summ2-8* are most similar to each other and *pat1-1path1-4path2-1summ2-8* positions in between *pat1-1path2-1summ2-8* and *pat1-1path1-4summ2-8*. Data File S1 exhibit differently expressed genes in multiple *pat* mutants which are clustered in Fig. 4B, and numbers of shared DE (differently expressed) genes are summed up in Fig. 4C. GO term analysis on misregulated genes in *pat1-1path1-4path2-1summ2-8* showed that: *i*. genes involved in oxidation-reduction are largely mis-regulated, *ii*. transcripts involved in oxidative stress response accumulated, and *iii*. transcripts responsible for auxin response and signaling and growth regulation are reduced in *pat* triple mutant (Fig. S2) (Huang et al., 2009).

**Fig 4.**
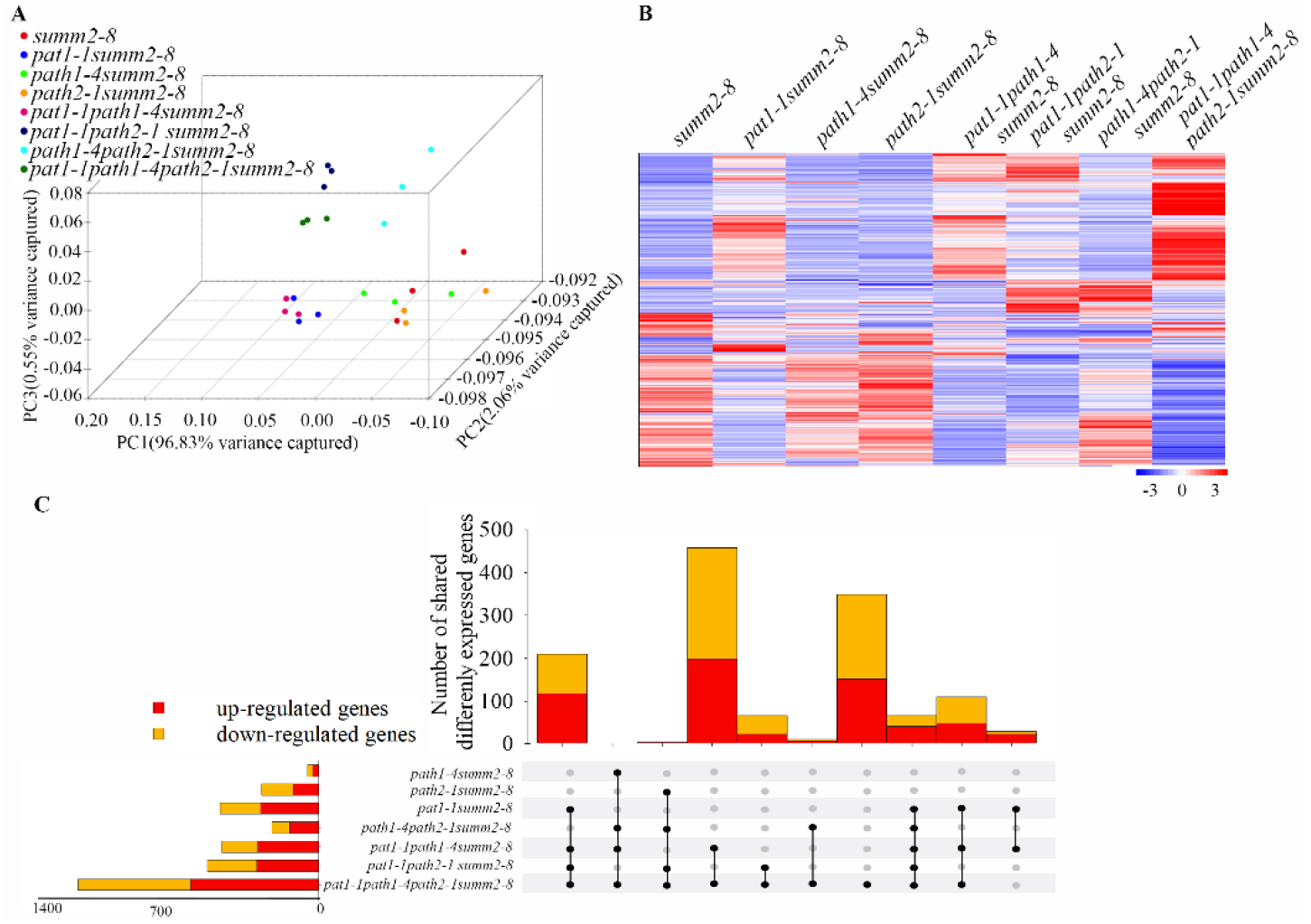
RNA-seq analysis on *pat* mutants. (**A**) 3-D PCA plot for general similarity of *pats* single, double and triple mutants with bio-triplicates. (**B**) Heat map clustering of differentially expressed genes for comparison of *pat* mutants and *summ2-8*. Pearson’s metrics were used in hierarchical clustering of the genes. In the plot, red indicates high expression and blue low expression. (**C**) Numbers of shared differently expressed genes in *pat* mutants.

### PATs regulate different genes expression during development

Leaf initiation and outgrowth have been reported under auxin regulation (Xiong, Y.&Jiao, Y, 2019). We therefore took closer look at the auxin responsive genes including *SAUR*s (*SMALL AUXIN UP RNA*s), *IAA*s, *MYB*s (*MYB DOMAIN PROTEIN*s), *WAG1, WAG2* etc. They were least expressed in *pat* triple mutant compared to all *pat* double mutants and *pat1* single mutant in which the auxin responsive genes were also down regulated compared to *summ2-8* and *path1* and *path2* single mutant (Fig. 5).

**Fig 5.**
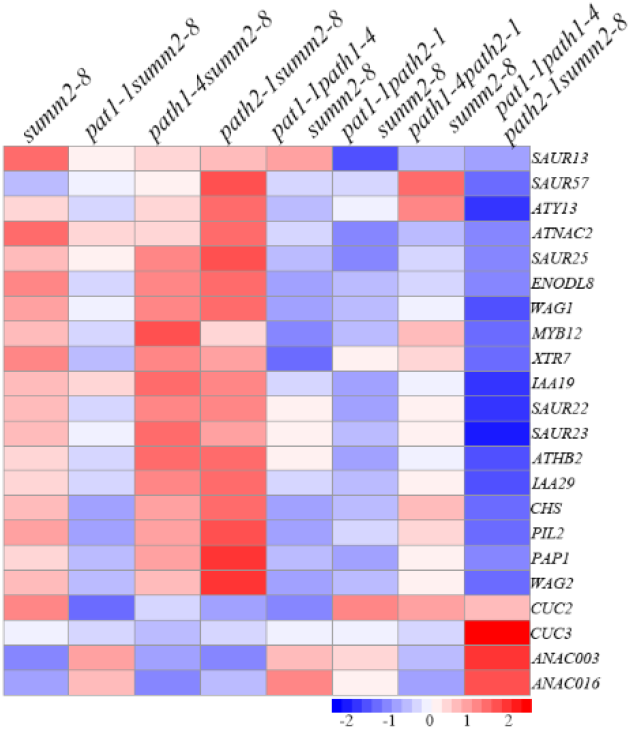
PATs regulate different genes expression. Heat map clustering of differentially expressed genes including auxin responsive genes, shade avoidance related genes and leaf serration related genes.

Light mediated petiole elongation during shade avoidance involves cell wall modification by XTHs (Xyloglucan Endotransglucosylase/Hydrolases) and is enhanced by PIFs (PHYTOCHROME-INTERACTING FACTORS) (Sasidharan, R., 2020; Xie, Y., 2017). *XTH7* and *PIL2/PIF6* were downregulated in *pat1-1summ2-8, pat1-1path1-4summ2-8* and *pat* triple mutants which exhibit shorter petioles than all other mutants (Fig.3& 5), suggesting PAT1 and PATH1 may regulate *XTH7* and *PIL2* to control petiole elongation.

The NAC family member CUC2 (CUP SHAPED COTYLEDON 2) and PIN1 auxin efflux carrier have been modeled to explain leaf serration and CUC3 has also been reported to be essential to keep serration (Bilsborough G.D., 2011; Serra L., 2020), however *CUC2* and *CUC3* were downregulated in the serrated mutant *pat1-1summ2-8* while we found other 2 NAC family gene *ANAC003* and *ANAC016* specifically accumulated in all serrated mutants indicating PAT1 and PATH1 may regulate other *NAC* genes expression for leaf patterning (Fig. 5). Nevertheless, all these results indicate all 3 PATs regulate different genes expression during leaf development.

## Discussion

Plant development requires massive overhauls of gene expression (Miyamoto et al., 2015). Our phenotypic characterizations showed that, compared to *pat1-1summ2-8*, leaf serration and dwarfism are more severe in *pat1-1path1-4summ2-8*, while *pat1-1path2-1summ2-8* exhibits less serration and normal petioles, indicating that PAT1 and PATH1 share some functions in leaf serration and petiole elongation while PATH2 may have opposite functions (Fig 3). However, loss-of-function of all 3 *PATs* causes severe dwarfism compared to any single or double mutants; together with this, *pat* triple mutant accumulates the most differently regulated genes of which 348 genes specifically misregulated in *pat* triple mutants (Fig 4C), indicating some redundancy and subfunctionaliztion among the three PATs in development (Duarte et al., 2006). The fact that all three PATs localized into PBs in response to IAA treatment also confirms functional redundancy of the PAT proteins (Fig. 2). While shared DE genes analysis manifests more DE genes shared by *pat1-1path1-4summ2-8* and *pat* triple mutant than the other two *pat* double mutants and *pat* triple mutant, probably suggesting mutation of *PAT1* and *PATH1* provide the highest contribution to the developmental phenotypes of *pat* triple mutant. Interestingly, *pat1-1summ2-8* mutants still exhibit dwarfism and leaf-serration, while neither *path1summ2-8* nor *path2summ2-8* mutants exhibit developmental defects (Fig1& 3). Our RNA-seq data shows that 382 transcripts are differentially expressed in *pat1-1summ2-8* compared to only 54 genes in *path1-4summ2-8* and 279 in *path2-1summ2-8* (Data File S1). While we do not know how these differences contribute to the developmental phenotype of *pat1*, these observations suggest a non-redundant function of PAT1 during plant development. In line with this, we previously reported that PAT1 specifically functions in response to osmotic stress (Zuo et.al., 2021) while all 3 PAT proteins function redundantly during *Turnip mosaic virus* infection (Zuo et.al., 2022) and other development processes including apical hook and lateral root formation (not published yet).

Besides *pat* tripe mutants, other mRNA decapping deficient mutants also exhibit abnormal development or postembryonic lethality (Xu et al., 2006; Xu and Chua, 2009; Perea-Resa et al., 2012; Zuo et al., 2022b). The stunted growth phenotype and downregulation of developmental and auxin responsive mRNAs in the *pat* triple mutant (Fig. 5&S2) may support a model in which defective clearance of suppressors of development hampers developmental programming. Several studies also implicated yeast PAT1-LSM1-7 complex inhibit exosome mediated 3’-5’ decay leading to certain mRNA stabilization and accumulation (Tharun, 2009; Gatica et al., 2019). Therefore, Arabidopsis *pats* loss-of-function mutants may also lose protection of some transcripts from exosome mediated decay, these destabilized transcripts could be auxin responsive genes and growth-related genes.

In conclusion, we have shown that PAT1, PATH1 and PATH2 interact with the LSM1 decapping factor and localize to PBs upon different stimuli perception (Fig. 1A &2). Through observing developmental phenotype of multi-*pat* mutants, investigating cellular localization of PAT proteins and performing RNA-seq to examine differently expressed genes we identified PAT1 serve as the main role during development especially in leaf morphology and work redundantly with other 2 PATs during plant development.

## Materials and Methods

### Plant materials and growth conditions

*Arabidopsis thaliana* ecotype Columbia (Col-0) was used as a control. All mutants used in this study are listed in Table S1. T-DNA insertion lines for At1g79090 (*PAT1*) *pat1-1* (Salk_040660), At1g12280 (*SUMM2*) *summ2-8* (SAIL_1152A06) and double mutant of *PAT1* and *SUMM2 pat1-1summ2-8* have been described (Petersen et al., 2000; Zhang et al., 2012; Roux et al., 2015). *pat1-1path1summ2-8* and *path2summ2-8* mutants were generated using CRISPR/CAS9 following standard procedures with plasmid pHEE401 containing an egg cell-specific promoter to control CRISPR/CAS9 (Wang et al., 2015). Independent T1 *pat1-1path1*^*het*^*summ2-8* and *path2*^*het*^*summ2-8* plants were used in crosses to produce *pat1-1*^*het*^*path1* ^*het*^*path2* ^*het*^*summ2-8*, Cas9 free mutants. Homozygous F2 plants were selected and used for experiments.

Plants were grown in 9×9cm or 4×5cm pots at 21°C with 8/16hrs light/dark regime, or on plates containing Murashige–Skoog (MS) salts medium (Duchefa), 1% sucrose and 1% agar with 16/8hr light/dark.

### Plant treatment and confocal microscopy

Seedlings were grown in liquid MS medium for 5 days, confocal microscopy pictures were taken with a Leica SP5 inverted microscope after 20min treatment with MS containing 0.5µM IAA, 1µM Cytokinin, 25µM ACC, 5µM flg22 or 0.1mM ATP.

### RNA-seq analysis

Total RNA was extracted from 6-week-old soil grown plants including *summ2-8, pat1-1summ2-8, path1-4summ2-8, path2-1summ2-8, pat1-1path1-4summ2-8, pat1-1path2-1summ2-8, path1-4path2-1summ2-8*, and *pat1-1path1-4path2-1summ2-8* using the NucleoSpin® RNA Plant kit (Machery-Nagel). RNA quality, library preparation and sequencing were performed by BGI. RNA-seq reads were mapped to the *Arabidopsis thaliana* TAIR10 reference genome with STAR (version 2.5.1b) using 2-pass alignment mode (Dobin et al., 2013). The read counts for each gene were detected using HTSeq (version 0.5.4p3) (Anders et al., 2015). The Araport11 annotation was used for both mapping and read counting. The counts were normalized using the TMM normalization from edgeR package in R (Robinson et al., 2010). PCA analysis were performed using IPython notebook (Wang and Ma’ayan, 2016). Prior to statistical testing the data was voom transformed and then the differential expression between the sample groups was calculated with limma package in R (Law et al., 2014; Ritchie et al., 2015). Genes with fold change ≥2 or ≤-2 and P-value ≤0.05 are listed in Data File S1. Functional Annotation Tool DAVID Bioinformatics Resources 6.8 has been used for GO term analysis for the differently expressed genes using detected genes as background (Huang et al., 2009).

### Cloning and transgenic lines

*PATs* promoter sequences with 5’ HindIII and 3’ XbaI linkers were amplified from Col-0 genomic DNA and cloned in plasmid pGWB515 to make pGWB515-PATsprom (Nakagawa et al., 2007). The Venus sequence without stop codon was amplified from pEN-L1-Venus-L2 (Mylle et al., 2013) and cloned in pGWB515-PATsprom. PAT genes were amplified from Col-0 genomic DNA and cloned into pENTR-D-TOPO (Invitrogen). The entry clones were combined with pGWB515-PATsprom and pGWB515-PATsprom-Venus to obtain N-terminal HA and Venus tags respectively (Nakagawa et al., 2007). pENTR-D-TOPO-GUS (Invitrogen) and pENTR-D-TOPO-LSM1 (Roux et al., 2015) were combined to pK7WGY2.0 (Karimi et al., 2007) to obtain N-terminal YFP tags. These fusions were transformed into *Agrobacterium tumefaciens* strain GV3101 for transient and stable expression. Transformants were selected on hygromycin (30 mg/L) MS agar, and survivors tested for transcript expression by qRT-PCR and protein expression by immuno-blotting. Primers for cloning are provided in Table S2.

### Transient expression, protein extraction and co-IP in *Nicotiana benthamiana*

*A. tumefaciens* strains carrying PAT fusions were grown in YEP medium supplemented with appropriate antibiotics overnight. Cultures were centrifuged and re-suspended in buffer (10mM MgCl2, 10 mM MES-K (pH 5.6), 100 μM acetosyringone) to OD600=0.8. *A. tumefaciens* strains carrying PATs-HA and YFP-LSM1 or YFP-GUS were mixed 1:1 and infiltrated into 3-week-old *N. benthamiana* leaves. Leaf samples for protein extraction and immunoprecipitation were collected 3 days post infiltration (dpi). Tissues for protein extraction were ground in liquid nitrogen and IP buffer (50mM Tris-HCl pH 7.5; 150 mM NaCl; 5 % (v/v) glycerol; 1 mM EDTA; 0.1%(v/v) NP40; 10 mM DTT; protease inhibitor cocktail (Roche); Phosstop (Roche)) added at 2mL/g tissue powder. Following 20 min centrifugation at 4°C and 13000 rpm, sample supernatants were adjusted to 2mg/ml protein and incubated 4 hours at 4°C with GFPTrap-A beads (Chromotek). Beads were washed 4 times with wash buffer (20 mM Tris pH 7.5; 150m M NaCl; 0.1 % (v/v) NP40 before adding 4×SDS buffer (novex)) and denatured by heating at 95°C for 5 min.

### Protein extraction, SDS-PAGE and immunoblotting

Tissue was ground in liquid nitrogen and 4×SDS buffer (novex) was added and heated at 95°C for 5 min, cooled to room temperature for 10min, samples were centrifuged 5min at 13000 rpm. Supernatants were separated on 10% SDS-PAGE gels, electroblotted to PVDF membrane (GE Healthcare), blocked in 5% (w/v) milk in TBS-Tween 20 (0.1%, v/v) and incubated 1hr to overnight with primary antibodies (anti-GFP (AMS Biotechnology 1:5.000, anti-HA 1:1,000 (Santa Cruz)). Membranes were washed 3 × 10 min in TBS-T (0.1%) before 1hr incubation in secondary antibodies (anti-rabbit or anti-mouse HRP or AP conjugate (Promega; 1: 5.000)). Chemiluminescent substrate (ECL Plus, Pierce) was applied before camera detection. For AP-conjugated primary antibodies, membranes were incubated in NBT/BCIP (Roche) until bands were visible.

## Figure

**Fig S1.**
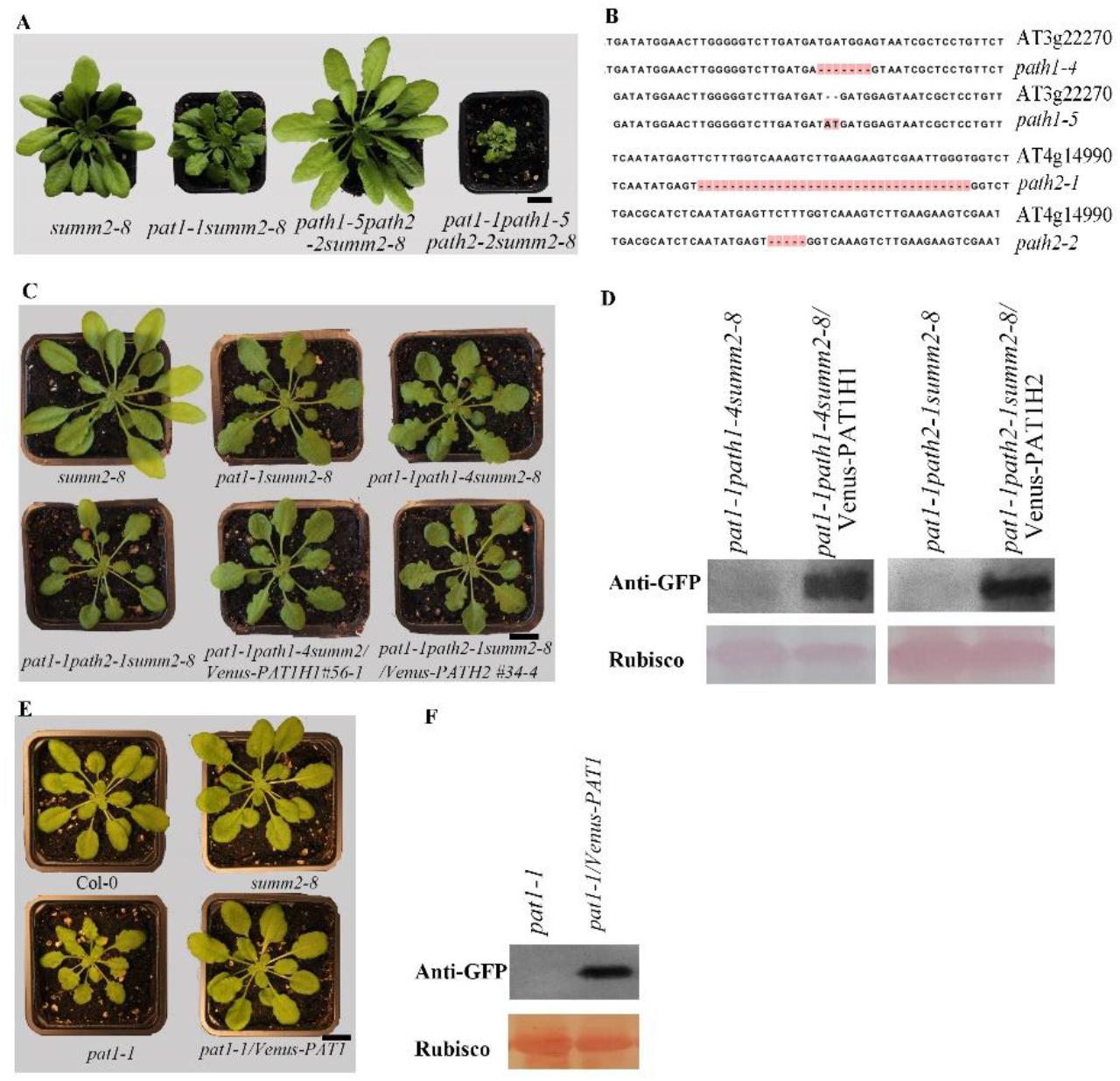
Characterization of *pat* mutants generated using CRISPR/CAS9 system. (**A**), 6 week-old plants of *summ2-8, pat1-1summ2-8, path1-5path2-2summ2-8* and *pat1-1path1-5path2-2summ2-8* grown in soil in chamber with 8/16hr light/dark at 21°C. One representative plant for each line is shown. The scale bar indicates 1cm. (**B**) sequence of independent *path1* and *path2* mutations. Western blots detecting the expression of PATH1 and PATH2 (**C**) and PAT1 (**E**) fusions with N-Venus and growth phenotype (**D&F**) of complemented lines. One representative plant for each line is shown. The scale bar indicates 1cm.

**Fig S2.**
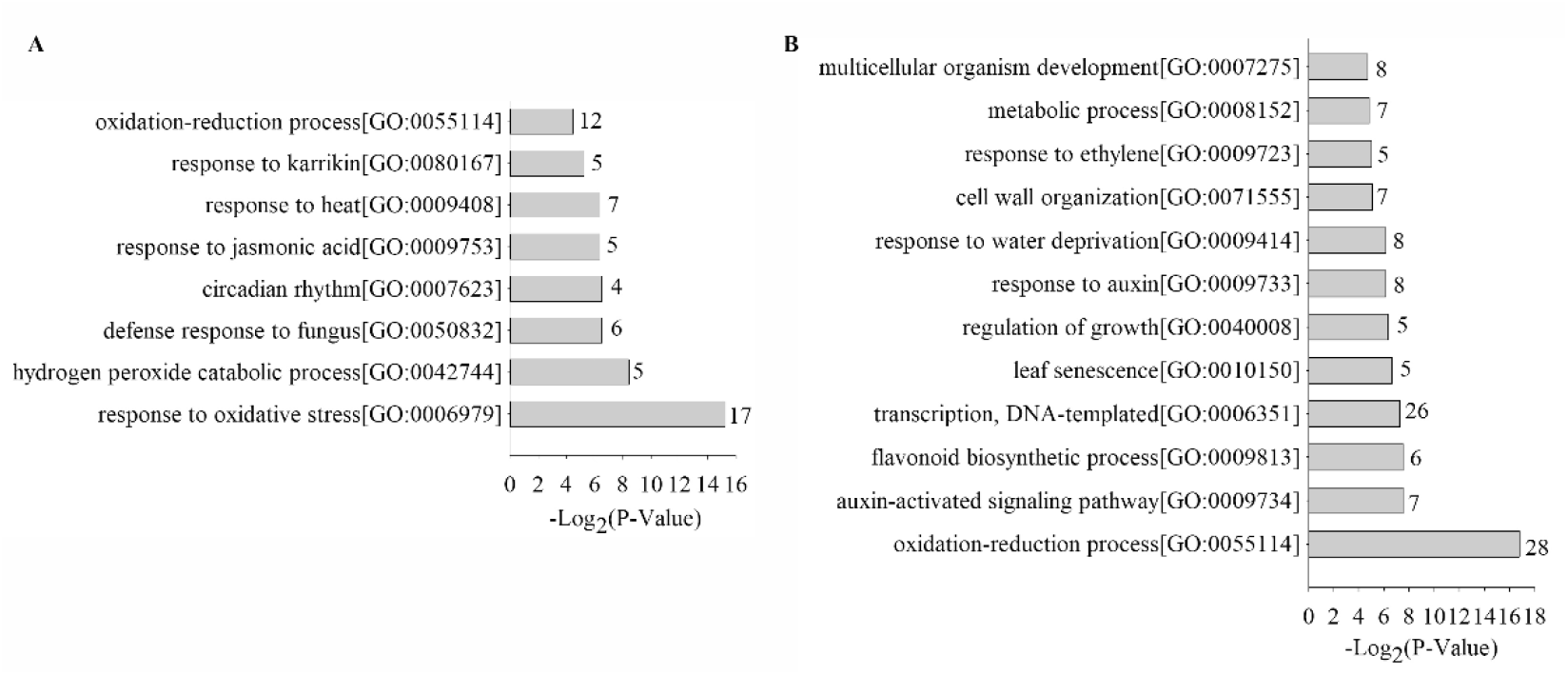
Analysis of differently expressed genes in *pat* triple mutants. Gene ontology analysis of transcripts upregulated (**A**) or downregulated (**B**) in 6-week-old plants of mRNA decay deficient mutant *pat1-1path1-4path2-1summ2-8* compared to *summ2-8*.

**Table S1.** Mutants used in this study.

| Mutant | Information | Source |
| --- | --- | --- |
| *path1-4* | 7bp deletion in Exon 2, frame shift and early stop codon | this study |
| *path1-5* | AT insertion in Exon 2, frame shift and early stop codon | this study |
| *path2-1* | 35bp deletion in Exon 2, frame shift, early stop codon | this study |
| *path2-2* | 5bp deletion in Exon 2, frame shift, early stop codon | this study |
| *pat1-1* | Salk_040660, T-DNA insertion in Exon 5 | Roux et al., 2015 |
| *summ2-8* | SAIL_1152A06, T-DNA insertion in Exon 1 | Zhang et al., 2012 |

**Table S2.**
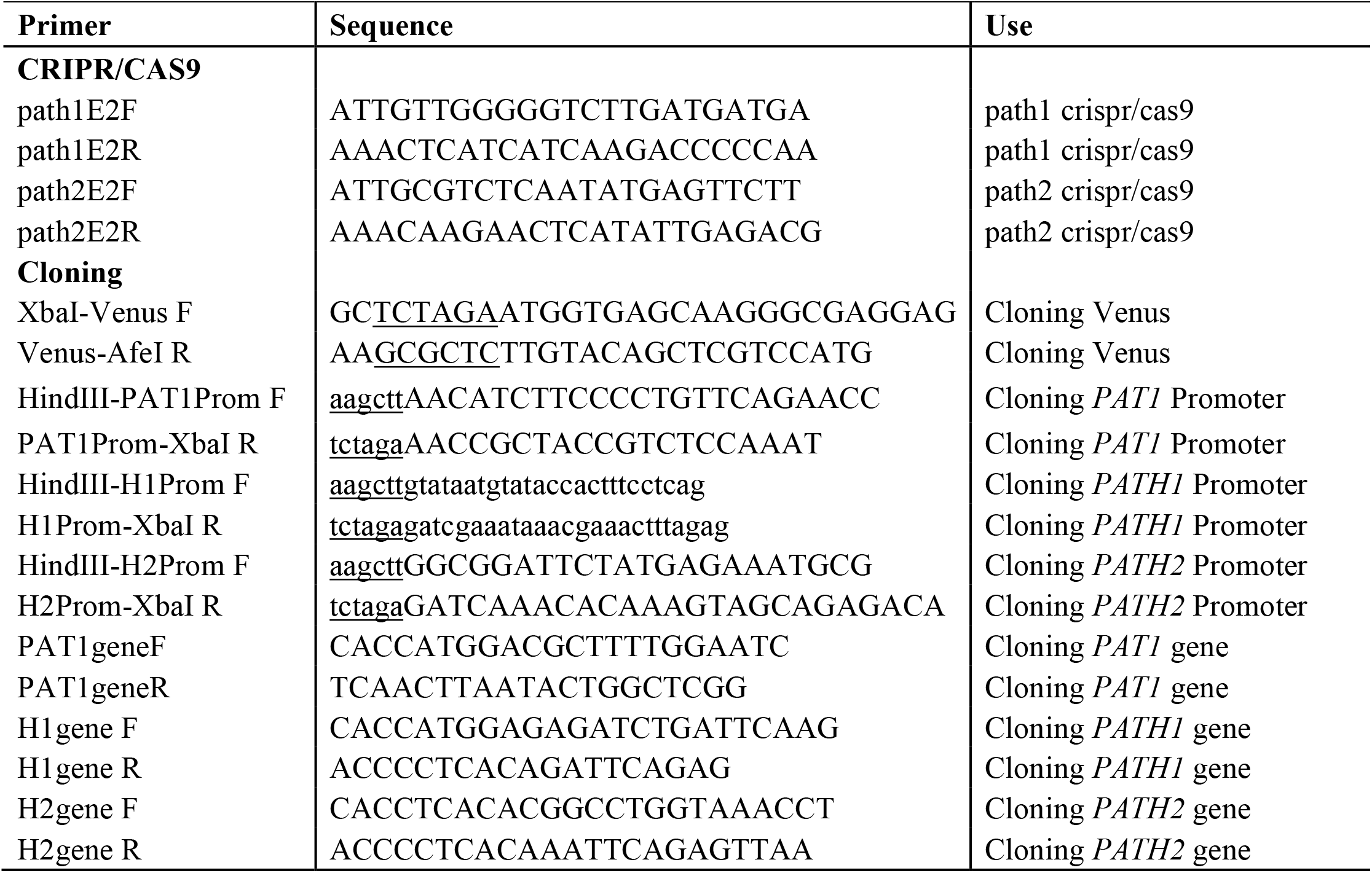
Primers used in this study.

## Acknowledgments

We thank Qi-Jun Chen for Phee401. Special thanks to John Mundy for advice throughout the project and critically reading the manuscript. We acknowledge the Bioinformatics and Scientific Computing Facility (VBCF) and Xiaoye Tong for the next-generation sequencing data analysis. This work was supported by the Novo Nordisk Fonden and the Hartmanns Fond to MP (NNF18OC0052967 and A32638) and a PhD scholarship from China Scholarship Council to ZZ (201504910714).

ZZ, MER, and MP conceived and designed the experiments. ZZ, MER, YD and ER performed experiments. ZZ and MP analyzed the data. ZZ and MP wrote the manuscript.

The authors declare no competing interests.

Correspondence and requests for materials should be addressed to MP.

